# Optimizing drug synergy prediction through categorical embeddings in Deep Neural Networks

**DOI:** 10.1101/2024.06.12.598611

**Authors:** Manuel González Lastre, Pablo González de Prado Salas, Raúl Guantes

## Abstract

Cancer treatments often lose effectiveness as tumors develop resistance to single–agent therapies. Combination treatments can overcome this limitation, but the overwhelming combinatorial space of drug-dose interactions makes exhaustive experimental testing impractical. Data-driven methods, such as Machine and deep learning, have emerged as promising tools to predict synergistic drug combinations. In this work, we systematically investigate the use of categorical embeddings within Deep Neural Networks to enhance drug synergy predictions. These learned and transferable encodings capture similarities between the elements of each category, demonstrating particular utility in scarce data scenarios.

## 1 Introduction

Molecularly targeted anticancer drugs have proven to be a form of therapy with remarkable results in recent years [1]. However, single agent treatments lose efficacy over time as tumours develop resistance to the drug administered [1, 2]. Moreover, single drugs are not always effective at the low doses needed to tolerate side effects. In this context, drug combinations have emerged as an alternative therapy with the potential to overcome resistance and increase efficacy at small doses [3].

In the quest for effective combination therapies, high-throughput preclinical screenings play an important role [4]. They allow experiments to be conducted in an undirected, parallelized manner, and constitute an invaluable tool for the massive interrogation of the potential effect of drug combinations. Drug combinations conferring synergy (usually quantified with respect to a null model where combined drug effects are additive or independent [5, 6]) are promising candidates for effective therapies. However, these empirical studies face several challenges, the main being the combinatorial explosion problem: the complete screening of *N* drugs and *D* doses needs *D*^*N*^ measurements, making exhaustive testing infeasible. Moreover, the mechanisms of action for many drugs remain poorly understood, further complicating the prediction of synergistic or antagonistic interactions between compounds [7].

To address the combinatorial complexity of drug synergy prediction, many efforts have been aimed at modeling the effects of compound combinations [8]. Previous efforts focused on mathematical and statistical models [9, 10, 11, 12, 13] that infer multifactorial responses from single-drug or pairwise data. Other strategies employed network methodology combined with biological and drug molecular information to predict drug interaction effects [14, 15]. More recently, machine learning (ML) techniques have been proven to be a useful tool in drug discovery [16, 17, 18], as they enable data-driven modeling without requiring prior mechanistic knowledge. In particular, the availability of large-scale molecular data—such as gene expression, somatic mutations, and drug structural descriptors—has facilitated the application of ML and deep learning (DL) methods to search for synergistic drug combinations [19, 20, 21, 22, 23, 24]. With some exceptions [21, 22], most of these approaches rely on high-dimensional omics and handcrafted drug features, making it difficult to tease apart the relevance of different features in synergy prediction and thus hindering interpretation.

Originally developed for unstructured data like text or images, embeddings have emerged as a transformative representation tool in ML. Rather than using sparse encodings (e.g., one-hot), embeddings map input categories into a dense, low-dimensional vector space that captures semantic similarity. These methods revolutionized natural language processing (NLP), starting with Word2Vec [25], which introduced dense word vectors and sparked the era of transfer learning in NLP. The benefits of embedding techniques are increasingly being applied to biological prediction tasks, where discrete, non-numeric entities such as drugs, targets, or cell lines must be represented in a meaningful, learnable form. Rather than relying on traditional one-hot or label encodings, several recent studies have shown that neural networks can learn dense representations of categorical biological inputs in an end-to-end manner to improve generalization and model performance. For instance, Liu et al.[26] combined sequence-based encodings with graph neural networks to embed drugs and targets in a heterogeneous biological network, learning task-specific representations of categorical nodes like proteins and compounds for accurate drug–target interaction prediction. Similarly, Li et al.[27] evaluated deep learning models that implicitly construct embeddings of drug and cell line identities using biological pathway information, demonstrating that these representations can enhance drug response prediction. These studies support the idea that embedding structured, categorical biological variables can uncover latent relationships and improve predictive performance— particularly in complex, data-sparse biomedical tasks. Categorical embeddings extend this principle to tabular data with high-cardinality variables. By training neural networks to learn task-optimized dense vectors for categorical inputs [28, 29], these methods can outperform traditional encodings and offer better scalability and transferability. Embedding-based representations thus hold particular promise for drug synergy prediction tasks, where categorical variables such as drug identity, mechanism of action, or cell line type encode rich relational structure that standard encodings fail to capture.

In this study, we systematically investigate the performance of different categorical embedding methods to predict the synergy of drug combinations. First, we train three different DL models specifically designed for tabular data, comparing both performance and training time required. Later, we use the learned embeddings as input features to the most widely used machine learning methods and compare their performance relative to traditional categorical encoding techniques. To evaluate these methods, we use, as a benchmark testing ground, the screening dataset provided in the AstraZeneca-Sanger Drug Combination Prediction DREAM Challenge [19], using the same synergy scores, data splits and performance metric used in the competition. In order to provide a controlled and focused assessment of the suitability of categorical embeddings for synergy prediction, we used a minimal set of input features for prediction containing both numerical (single drug response features) and categorical variables, without further use of molecular or drug attributes. We show that for 10 out of the 14 benchmarked models, the learned embeddings outperform or tie with traditional encodings and in 5/14, the performance increases by more than 30%.

## 2 Materials and Methods

### 2.1 The dataset

We based our analyses on data from Sub-challenge 1 (SC1) of the AstraZeneca-Sanger Drug Combination Prediction DREAM Challenge [19]. This challenge provided a large-scale in vitro dataset, comprising 3,475 experiments involving 167 unique drug combinations tested across 85 cancer cell lines. Each experiment measured the viability of a cell line treated with a particular drug pair, from which a single synergy score was derived using the Loewe model of additive combinations[30, 19]. Positive synergy scores indicate that the combined effect is greater than expected from additivity, zero corresponds to an additive combination, and negative values indicate antagonism.

The cancer cell lines span six different cancer types, predominantly breast (34 cell lines), lung (22), bladder (14), and gastrointestinal (12) cancers. Each cell line is characterized by features such as the cell line name, tissue type (`GDSC tissue descriptor 2`), microsatellite instability status (`MSI`), and growth properties (adherent or suspension culture).

The drug combinations consist of pairs drawn from 118 unique compounds, including both targeted therapies and chemotherapeutic agents. For each compound (referred to as `Compound A` and `Compound B` of the pairwise combination), the dataset provides detailed information, including putative targets (`Putative target_yA` and `Putative target_B`), general functional annotations based on the dominant biological process attributed to the target(`Function_A`, and `Function_B`) and associated signaling pathways (`Pathway_A` and `Pathway_B`) (Figure 1B). Single–drug dose–response properties are also included for each compound (Figure 1A), such as the maximum tested concentration (`Max. conc. A` and `Max. conc. B`), half-maximal inhibitory concentration (IC_50_ A and IC_50_ B), Hill coefficient (`H A` and `H B`), and maximal effect (E_inf_ A and E_inf_ B).

**Figure 1.**
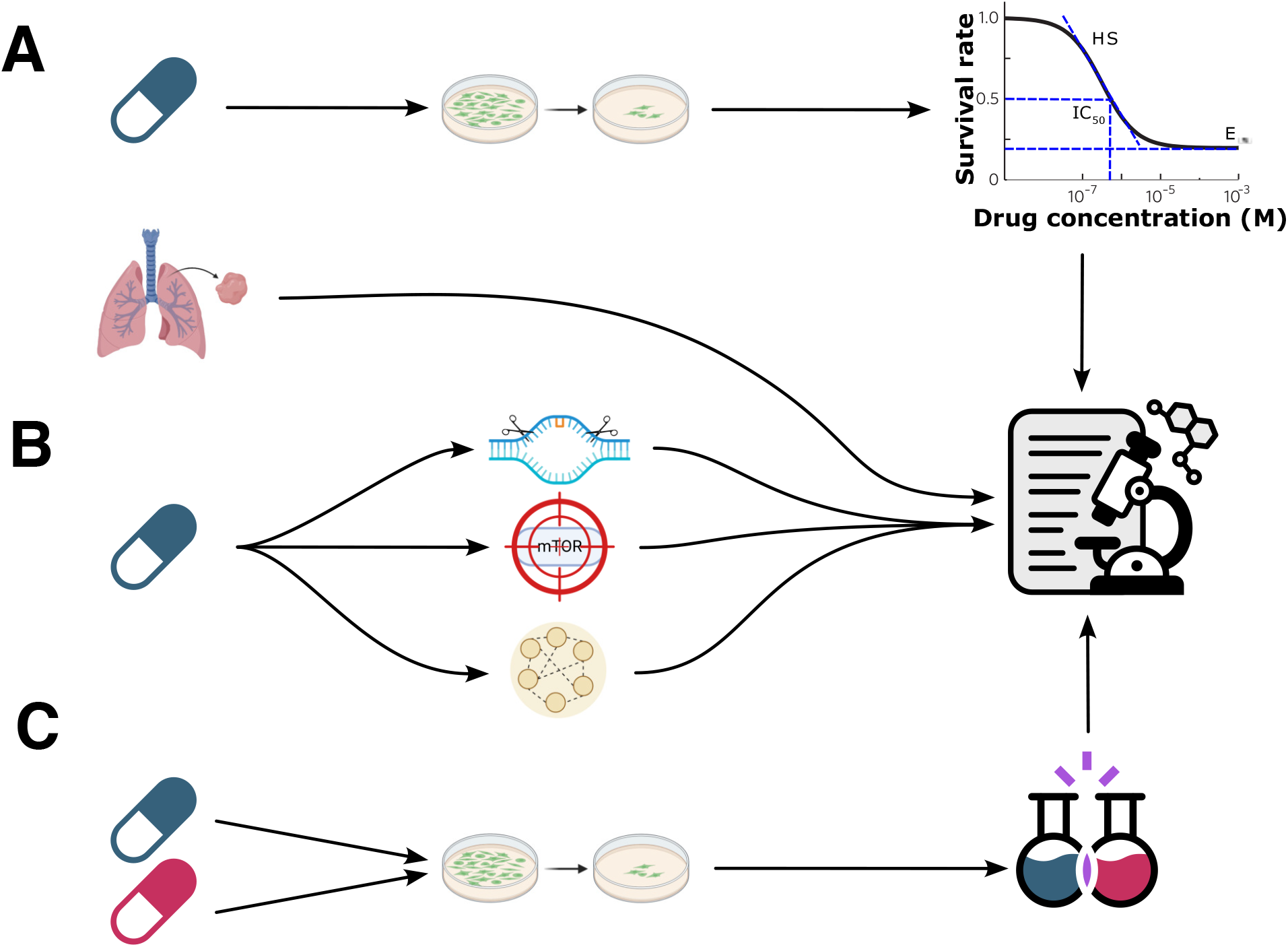
Input features and output target for synergy prediction. The dataset of input features for model training consists of: **A**. Numerical variables, corresponding to parameters of the single drug experimental dose-response curve: half-maximal inhibitory concentration (IC_50_), slope of response (Hill coefficient H S) and drug efficacy (maximal effect E_inf_), as well as maximum tested concentration. **B**. Categorical variables: cell line and tissue descriptor, and drug attributes (function, molecular target and pathway) as well as drug names. As output for training and prediction, we use the experimentally measured synergy scores of pairwise drug combinations (**C**).

Our analysis focuses on utilizing the available pharmacological and cell line features without incorporating other biological and high-throughput data, such as copy-number variation, DNA methylation, transcriptomic profiles, etc. The goal is to predict the synergy score of drug combinations based on this minimal set of features (Figure 1C), to assess in a systematic and controlled manner the improvement in predictive performance achieved when using categorical embeddings, relative to more traditional encoding methods.

### 2.2 Data pre–processing and feature selection

We conducted several pre–processing steps to prepare the data (available as Supplementary Data 1 in the original DreamChallenge competition paper[19]) for model training:

1. Only high quality experiments (flagged as `QA=1`, reliable measurements) were retained.
2. Drug synergy data was enriched with cell line and drug portfolio information available in the original dataset. For cell lines, we incorporated the following categorical features: `GDSC tissue descriptor 2, MSI`(microsatellite instability), and cell `Growth properties`. For the drug portfolio, we included the `Putative target`, Function and `Pathway` for each drug.

Feature selection involved dropping unnecessary or redundant columns, including `Challenge, QA, COSMIC ID`, and columns with over 20% missing values such as H-bond acceptors (`HBA`), H-bond donors (`HBD`), `Molecular weight, cLogP, Lipinski`, and `SMILES`. Additionally, `GDSC tissue descriptor 1` and `TCGA label` were removed due to redundancy with `GDSC tissue descriptor 2`. The remaining features were grouped into categorical features: (`Cell line name, Compound names, GDSC tissue descriptor 2, MSI, Growth properties, Putative target, Function` and `Pathway`) of drugs; and numerical features (monotherapy dose-response parameters for each single drug as described above). The final dataset consisted of **12 categorical features and 8 numerical features**, with the synergy score as the target variable.

After pre–processing, the dataset was divided into training, test, and validation(LeaderBoard) sets, following the original splits from the DREAM Challenge Subchallenge 1, where the objective task is to predict the synergy score for drug combinations for which training data on those same combinations are available. From a total of 3,475 experimental pairwise combinations, 1,795 combinations were used for training, 1,089 for test and the rest (591) were kept for prediction (LeaderBoard set).

### 2.3 Evaluation metric

Due to significant experimental variability in high-throughput screenings, traditional regression metrics like mean squared error are less effective for evaluating synergy predictions. We adopted the Weighted Pearson Correlation (WPC) coefficient, as proposed in the original DREAM Challenge [19], to assess model performance. The WPC is defined as:

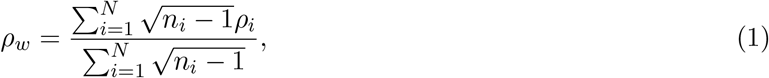

where *ρ*_*i*_ is the Pearson correlation coefficient between experimental and predicted synergy scores for drug combination *i, n*_*i*_ is the number of cell lines tested with drug combination *i*, and *N* is the total number of drug combinations. This metric emphasizes correlation across different cell lines rather than precise prediction of synergy scores. All WPC values reported in this study are calculated using the LeaderBoard (LB) dataset of the original competition, comprising 591 drug combinations for prediction. As a reference, replicate experiments in the original screening protocol achieve average WPC values around 0.4 [19].

### 2.4 Encoding Methods

#### 2.4.1 Traditional encoding methods

We employed traditional encoding techniques such as Label Encoding and One-Hot Encoding. Label Encoding is the simplest form of encoding and consists of assigning a unique integer to each category, preserving dimensionality but potentially introducing unintended ordinal relationships between different items of the category. On the other hand, One-Hot Encoding transforms each categorical variable into binary columns representing the presence or absence of each category, which does not introduce any ordinal bias at the expense of increasing the dimensionality.

#### 2.4.2 Embedding encodings

Categorical embeddings are proposed as a solution to the trade-off between bias and dimensionality that traditional encodings present. Inspired by techniques from Natural Language Processing [25], categorical embeddings map item classes inside a categorical variable to dense vectors in a continuous space. Since the dimensionality of the output’s vector space is smaller than the dimensionality of the input’s, it forces the neural network to abstract properties from the elements inside a category.

Embeddings enable the model to learn and transfer knowledge between the categorical variables (drug and cell line information) based on the resulting network output (synergy score). The captured relationships between items of categorical variables could thus enhance predictions, specially in biological applications where data are scarce. To train and evaluate the models, we use the PyTorch Tabular library [31], which implements state–of–the–art tabular learning models, abstracts training through its *TabularModel* class and later facilitates extracting the learned embeddings with the **scikit–learn** compatible *EmbeddingTransformer* class.

We train three of the models implemented in the library: the CategoryEmbedding model, which is the simplest one and consists of a feedforward neural network utilizing embedding layers for categorical variables[29]; AutoInt (Automatic Feature Interaction Learning via Self-Attentive Neural Networks) [32], which utilizes multihead self–attention and an interacting layer to promote interactions between different features; and TabTransformer [33], which incorporates Transformer-based attention mechanisms to learn dependencies between features. We maintain the same hyperparameters for all three models across training runs: batch size of 512, a maximum of 200 epochs, and early stopping patience of 5 epochs.

### 2.5 Benchmark models selection

Once the embedding vectors for our categorical features have been pre-trained with the three methods outlined above, we use them as inputs for a wide range of machine learning models commonly used for tabular data, categorized by their nature as follows:

- **Linear models:** Linear Regression, Ridge Regression, Lasso Regression, ElasticNet Regression, Bayesian Ridge Regression, and Stochastic Gradient Descent (SGD) Regressor.
- **Kernel-based and non-linear models:** Support Vector Regressor (SVR) and Gaussian Process Regressor.
- **Tree-based models:** Decision Tree Regressor, Random Forest Regressor, and Extra Trees Regressor. Additionally, boosting methods were employed, given their iterative improvement of weak learners. These models included AdaBoost Regressor, Gradient Boosting Regressor, and XGBoost Regressor

We use the **scikit–learn** [34] implementation for all models, except for XGBoost for which we employ the *dmlc/xgboost* [35] library.

To assess the performance of different embedding techniques, we also compare them with traditional label and One-Hot Encoding methods across the whole range of machine learning models. The computational pipeline is sketched in Figure 2.

**Figure 2.**
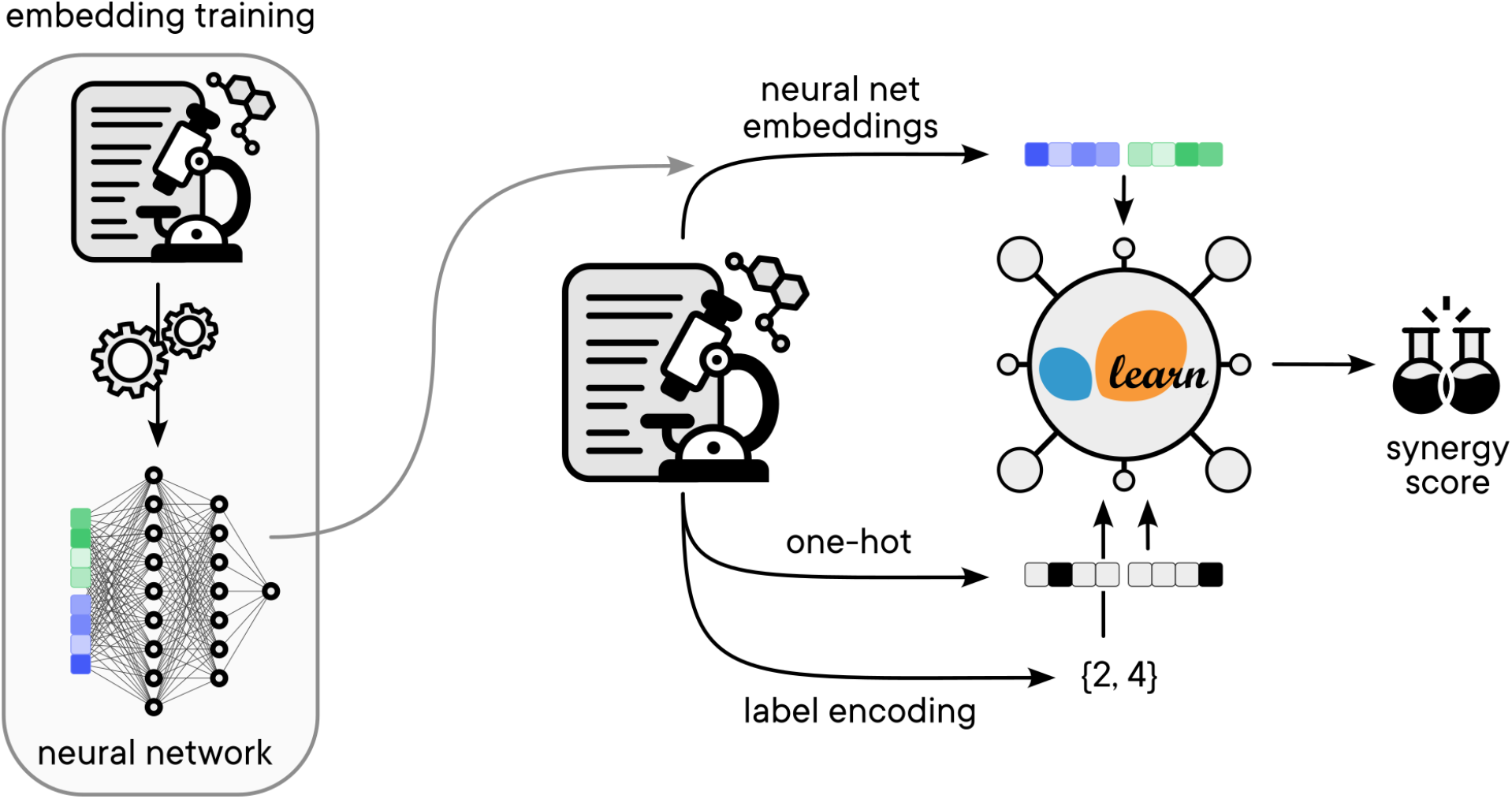
Training and prediction pipeline. The processed dataset is used to train deep learning models. First, we optimize categorical embeddings for drug synergy prediction (left). These learned embeddings are subsequently employed as input features for scikit-learn models, together with the numerical features outlined in Methods. The final performance of the pipeline is benchmarked against traditional encoding methods, such as label encoding and One-Hot Encoding.

### 2.6 Dimensionality reduction and clustering

To visualize relationships between the embeddings of items of a given category we used two well established methods for non-linear dimensionality reduction: the t-Distributed Stochastic Neighbour Embedding (t-SNE)[36] and the Uniform Manifold Approximation and Projection (UMAP)[37]. t-SNE analyses were done with the R package *Rtsne* with the perplexity parameter (number of close neighbors) set equal to 3. For UMAP, we used the R package *uwot* with 3 nearest neighbors and default settings. Next, we applied k-means clustering to group items within each category.

To compare clusterings in the plane of two main components for different embedding and dimensionality reduction methods, we calculated the adjusted rand index (relative proportion of element pairs that belong to the same cluster, corrected for random chance coincidences). For a better visual inspection, in Figure 3 and Figure S1 we calculated the overlap (number of common elements) between all cluster pairs of two different clusterings, and assigned the same colors to those clusters with maximum overlap. For Figure 3D-F (distributions of experimental drug combinations corresponding to different Compound B clusters) we calculated how many combinations were shared among the same clusters of different embedding methods (normalized by the smaller cluster size), as a measure of relative overlap of the distributions.

**Figure 3.**
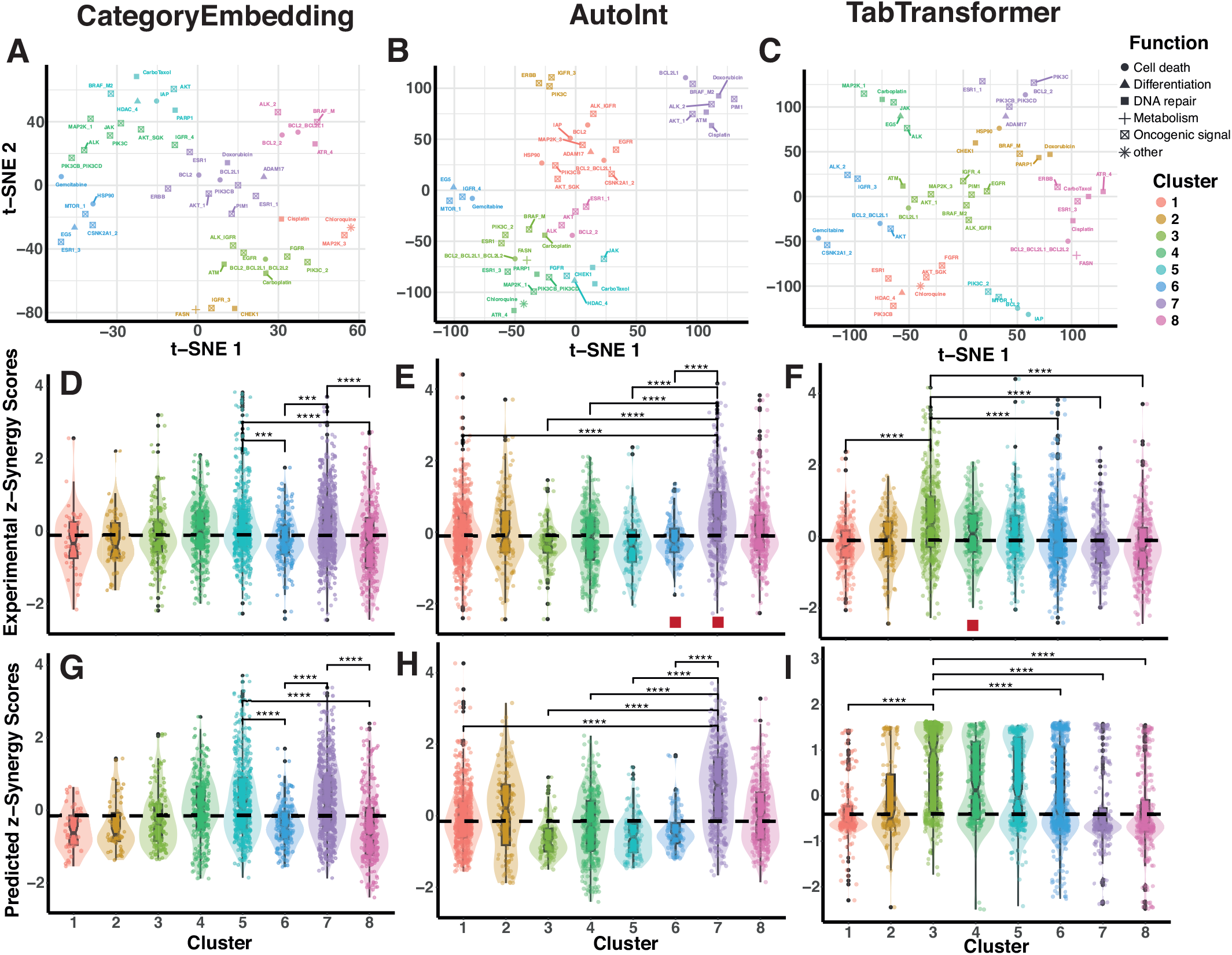
Differences in embedding vector information among methods. Analysis of embedding vectors for the categorical variable Compound B, containing 47 different drug names (most of them associated to their putative molecular targets, labels in top panels). **A-C**. Embedding projections on the first two principal dimensions of t-distributed stochastic network embedding(t-SNE) [36] for the three embedding methods tested. Colors correspond to different clusters using a k-means clustering algorithm after dimensionality reduction. In AutoInt and TabTransformer, we assigned the same colors to clusters with maximum overlap of items with CategoryEmbedding. Symbols denote the dominant functionality associated to the drug target, as grouped in the original dataset [19]. **D-F**. Boxplots with overlaid distributions (violin plots) of the z-normalized experimental synergy scores corresponding to the drug combinations containing Compound B, grouped by the clusters shown in the top panels. Shown with bars are the most significant differences between distribution means. Stars represent adjusted (Bonferroni) p values calculated with two-sided Wilcoxon rank-sum tests (ns: p *>* 0.05; *: p ≤ 0.05, **: p ≤ 0.01; ***: p ≤ 0.001; ****: p ≤ 0.0001). The dashed horizontal line represents the median value of the whole distribution of synergy scores. Red squares in **G** and **F** indicate clusters sharing more than 70% experimental combinations with the corresponding clusters in **E. G-I**. Boxplots of the predicted z-normalized synergy scores for the same drug combinations shown in the middle panels.

## 3 Results and discussion

### 3.1 Performance of PyTorch Tabular Models

First, we evaluate the performance and computational cost of the PyTorch Tabular models for predicting drug synergy scores. We train three models—AutoInt, CategoryEmbedding and TabTransformer—using the dataset described in the Methods section, and compute the Weighted Pearson Correlation (WPC) for each model as a measure of prediction accuracy. Table 1 shows that, on average, AutoInt achieves the highest accuracy amongst the three, with a WPC of 0.270. In contrast, CategoryEmbedding is the fastest model, requiring an average of 4 seconds per training run but achieving a lower WPC of 0.184. TabTransformer strikes a middle ground with a WPC of 0.225, though it is the slowest to train, averaging over 9 seconds per run.

**Table 1:** Performance and training time of PyTorch Tabular models. The table summarizes the Weighted Pearson Correlation (WPC) and training times of three PyTorch Tabular models: CategoryEmbedding, AutoInt and TabTransformer. Training was performed using an NVIDIA RTX™ A1000 GPU. Results are averaged over 10 random seeds and reported as mean (standard deviation).

| Model | WPC | Training time (s) |
| --- | --- | --- |
| CategoryEmbedding | 0.184 (0.031) | **3.996 (0.264)** |
| AutoInt | **0.270 (0.015)** | 8.267 (0.118) |
| TabTransformer | 0.225 (0.020) | 9.368 (0.088) |

### 3.2 Analysis of different embedding models

To gain some insight into the relationships between categorical features captured by the different embedding models, we performed a systematic analysis of similarities between the items in a given category of drug information(`Compound, Putative target, Pathway, Function`) using dimensionality reduction methods, see section 2.6 in *Materials and Methods*.

As an example, we show in Figure 3A-C the t-SNE embedding projections of the category Compound B, for the three different methods employed in this work: CategoryEmbedding(CE), AutoInt(AI) and TabTransformer(TT). For a controlled comparison, these embeddings correspond to trainings of the three methods initialized with the same random seed, providing a performance close to the averages shown in Table 1. Several groups of Compound B items (labels in the plots) are well separated in the two main t-SNE dimensions. However, most of the clusters have a small overlap of items among the different embedding methods (adjusted rand indices 0.02 for CE vs. AI embeddings, 0.01 for CE vs. TT, and 0.04 for AI vs. TT). These differences in cluster composition are not caused by the specific dimensionality reduction technique or random seed, as cluster overlap for the same embedding method using t-SNE or UMAP projections is very high (adjusted rand indices between 0.54-0.73, see Figure S1). The small cluster overlap among embeddings rather indicates that the three embedding methods learn different relationships between variables and items. We also notice that clusters do not seem to be associated to a specific functionality of the drug targets (coded by the different symbols in Figure 3A-C).

To visualize more explicitly the differences between the three embedding methods, we collect the experimental synergy scores of the training dataset and group them by their corresponding Compound B labels associated to each cluster, Figure 3D-F. We see that many of the clusters revealed by the embeddings display statistically significant differences among synergy scores (we show the top 4-5 most significant differences in the plots). For example, clusters 6 and 8 in CategoryEmbedding contain synergy scores significantly below the median value, while clusters 5 and 7 include significantly higher scores. Figure 3 highlights the fact that the embedding vectors code some useful relationships among items that allow to capture trends in the data. However, relationships and trends differ among embedding methods, as suggested by the small overlap between clusters. Still, there are a few clusters that are consistently reproduced among methods, marked with red squares in Figure 3E-F (see Figure caption).

To see how the predictions align with these trends in the experimental data, we show in Figure 3 G-I the predicted synergy scores grouped by the same clusters. Notably, the trends in experimental data are not only enhanced in the predictions (confirming that the embeddings capture the visualized trends in data), but reveal the different ways in which these trends are encoded in the embedding. The simplest method, CategoryEmbedding, shows the best overall and by-cluster correlations between experimental and predicted trends (Supplementary Figure S2). In AutoInt, predictions overemphasize the experimental trends. With this method, the biases observed in some clusters, for instance in clusters 3, 5 and 6 toward low experimental synergy scores, are pushed to lower predicted values (notice the overlaid distributions in the plots). Finally, TabTransformer reveals a clear separation between low and high synergy score values, with most of the clusters showing skewed bimodal distributions of predicted synergy scores. These differences in data biases can also be observed in the correlations between experimental and predicted scores, Figure S2. From the three Neural Network encodings tested, TabTransformer performs worse on individual synergy predictions, as it tends to separate high and low synergy score values as much as possible.

### 3.3 Performance of encoding methods across machine learning models

Finally, we compare the prediction accuracy of the learned embedding encodings (AutoInt, CategoryEmbedding, and TabTransformer) against two standard methods, Label Encoding and One-Hot Encoding, by training a broad selection of machine learning models. As shown in Table 2, each model was evaluated over ten random initializations to ensure that the results are statistically significant. The Table includes both the average and the standard deviation to show that the training is robust across different seeds.

**Table 2:**
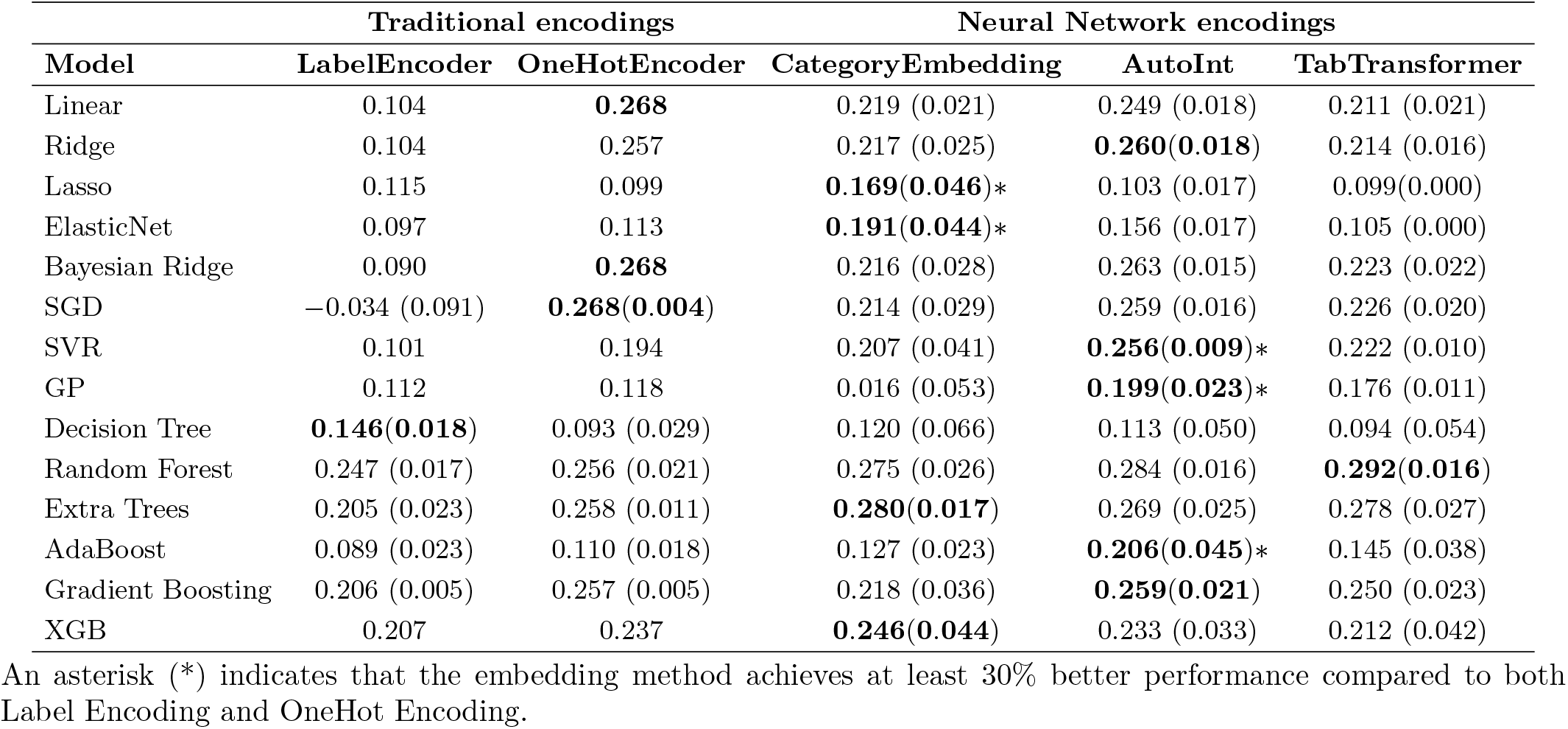
Performance summary (Leaderboard subset) of traditional encoding methods versus learned neural network embeddings across various models. Results are averaged over 10 random seeds and reported as mean (standard deviation). Standard deviation is omitted for deterministic training runs.

| Model | Traditional encodings |  | Neural Network encodings |  |  |
| --- | --- | --- | --- | --- | --- |
|  | LabelEncoder | OneHotEncoder | CategoryEmbedding | AutoInt | TabTransformer |
| Linear | 0.104 | **0.268** | 0.219 (0.021) | 0.249 (0.018) | 0.211 (0.021) |
| Ridge | 0.104 | 0.257 | 0.217 (0.025) | **0.260(0.018)** | 0.214 (0.016) |
| Lasso | 0.115 | 0.099 | **0.169(0.046)\*** | 0.103 (0.017) | 0.099(0.000) |
| ElasticNet | 0.097 | 0.113 | **0.191(0.044)\*** | 0.156 (0.017) | 0.105 (0.000) |
| Bayesian Ridge | 0.090 | **0.268** | 0.216 (0.028) | 0.263 (0.015) | 0.223 (0.022) |
| SGD | −0.034 (0.091) | **0.268(0.004)** | 0.214 (0.029) | 0.259 (0.016) | 0.226 (0.020) |
| SVR | 0.101 | 0.194 | 0.207 (0.041) | **0.256(0.009)\*** | 0.222 (0.010) |
| GP | 0.112 | 0.118 | 0.016 (0.053) | **0.199(0.023)\*** | 0.176 (0.011) |
| Decision Tree | **0.146(0.018)** | 0.093 (0.029) | 0.120 (0.066) | 0.113 (0.050) | 0.094 (0.054) |
| Random Forest | 0.247 (0.017) | 0.256 (0.021) | 0.275 (0.026) | 0.284 (0.016) | **0.292(0.016)** |
| Extra Trees | 0.205 (0.023) | 0.258 (0.011) | **0.280(0.017)** | 0.269 (0.025) | 0.278 (0.027) |
| AdaBoost | 0.089 (0.023) | 0.110 (0.018) | 0.127 (0.023) | **0.206(0.045)\*** | 0.145 (0.038) |
| Gradient Boosting | 0.206 (0.005) | 0.257 (0.005) | 0.218 (0.036) | **0.259(0.021)** | 0.250 (0.023) |
| XGB | 0.207 | 0.237 | **0.246(0.044)** | 0.233 (0.033) | 0.212 (0.042) |

An asterisk (\*) indicates that the embedding method achieves at least 30% better performance compared to both Label Encoding and OneHot Encoding.

For 10 of the 14 benchmarked models, embeddings either outperform or at least tie with traditional encoding methods and in 5 of them, the improvement was more than 30%. This trend holds consistently across multiple random initializations, as shown in Tables S1 and S2, where we analyze performance variability due to changes in the embedding and *sklearn* random seeds separately. Embeddings demonstrate a clear advantage in tree-based models, where they consistently outperform Label and One-Hot Encoding, with Decision Tree being the only exception. The same can be said of the kernel–based models, Support Vector Regressor and Gaussian Process, where embeddings show substantial improvements of 32% and 68% respectively (Table S3). The models with the largest observed improvements are ElasticNet (+70%) and AdaBoost (+88%) (Table S3).

When comparing the three embedding approaches, differences emerge in their effectiveness across models. CategoryEmbedding improves over traditional encodings in 7 out of 14 models, AutoInt for 8/14 and TabTransformer only for 5/14. Moreover, CategoryEmbedding and AutoInt are the best performing encoding in 4/14 and 5/14 models respectively, while TabTransformer is only the best performing method for the Random Forest model (which represents the best performing model–embedding pair in our benchmark).

These differences in performance can be better understood by analyzing the synergy score distributions across clusters represented in the embeddings, as illustrated in Figure 3G-I. Both CategoryEmbedding and AutoInt replicate more consistently the experimental synergy score distributions within their respective clusters. In contrast, TabTransformer does not reproduce these distributions accurately, exaggerating biases as seen in the skewed or bimodal predicted score distributions in Figure 3I. These discrepancies likely contribute to TabTransformer’s more inconsistent performance across models, with the exception of the Random Forest model.

So far, the focus of this work has been to systematically determine the role of categorical embeddings in improving drug synergy predictions at the single-model level. However, competitive machine learning pipelines usually utilize some form of ensembling. To explore the benefit of using ensembles of different models trained with categorical embeddings, we adopted two simple ensembling strategies based on greedy forward selection [38] (Supplementary Information section S3). Using as a starting point the embeddings of a model with a relatively good performance, our results show a consistent improvement up to WPC 0.32-0.33. These results approach the experimental performance average attained by biological replicates (WPC ∼ 0.4) [19]), despite being trained on a minimal set of features. This highlights the strength of the learned embeddings in capturing relevant pharmacological and cellular information.

## 4 Conclusions

In this work, we tested the use of categorical embeddings within deep neural networks for drug synergy prediction, demonstrating their advantages over traditional encoding strategies such as Label Encoding and One-Hot Encoding. Using a minimal set of input features—focusing solely on pharmacological and cell-line properties—we systematically trained three neural network models designed for tabular data and examined the embedding representations they learned.

Our analyses showed that these embeddings capture relationships between labels included in cell-line and drug categorical variables, revealing trends in the data based on this information. This indicates that embeddings can encode meaningful patterns within categorical data for our particular task, which traditional encoding techniques fail to capture. However, there seems to be marked differences in the type of relationships embodied by the different methods. This is highlighted both by the different clustering of drug combinations based on dimensionality reduction of the embedding vectors, and by the systematic biases in the predicted scores. These observations indicate that there is no ‘one-size-fits-all’ recipe for improving synergy prediction, particularly when embeddings are combined with machine learning models of different nature.

In this sense, our results demonstrate that learned embeddings either outperform or tie with traditional encoding methods in 10 out of the 14 benchmarked machine learning models, with performance improvements exceeding 30% in 5 cases. Notably, embeddings showed clear advantages in tree-based and kernel-based models, including Random Forest, Extra Trees, Support Vector Regressor, and Gaussian Process Regressor, with the latter two achieving improvements of 32% and 68%, respectively. Among the three embedding methods evaluated, AutoInt and CategoryEmbedding demonstrated broader success across models compared to TabTransformer, suggesting that explicit feature interaction mechanisms in AutoInt and the simplicity of CategoryEmbedding may generalize better in low-feature biological settings.

To place the results shown here into context, the best performing methods in Table 2 are slightly better than the average performance across all teams of the original DreamChallenge competition (WPC = 0.24 *±* 0.01, weighted Pearson correlation *±* standard error). Most of the teams, however, used deep molecular information of cell lines (gene expression, somatic mutations, DNA methylation and copy-number alterations) as well as extensive feature engineering. When ensembling results of our categorical embeddings across different models, our overall performance is comparable to that of one of the winning teams (NAD) including cell molecular features and detailed drug annotation (see Fig. 4 in Menden et al.[19]).

We emphasize that, for consistency, we used the same overall performance index (WPC) and synergy score quantification employed in the original DreamChallenge competition. Other definitions and methods to quantify synergy may represent more accurately the experimental effect of drug combinations, or improve predictions in data-driven approaches[39, 40, 41, 42, 21]. In the future, it would be interesting to explore the performance of these embedding methods with alternative synergy indicators, as well as in different drug screenings. Moreover, we did not use available high-throughput and drug molecular information for training and prediction. We foresee that the use of this kind of biological information, combined with categorical embeddings, may significantly advance both the predictive power and the interpretability of Machine and Deep Learning tools for this task.

In summary, our study establishes categorical embeddings as a scalable and data-efficient alternative to traditional encoding strategies for drug synergy prediction. These embeddings could be of particular advantage in biological datasets where experiments are scarce, as they can be reused across different tasks enabling learning transferability.

## Acknowledgments

This work was supported by the Spanish Ministry of Science and Innovation and the “ERDF A Way of Making Europe” program. M. G. L. acknowledges financial support through project PID2020-115864RB-I00 and grant PRE2021-098697. R. G. acknowledges support from project PID2021-122936NB-I00. Both M. G. L. and R. G. acknowledge additional support from the “María de Maeztu” Programme for Units of Excellence in R&D (CEX2023-001316-M).

## Author contributions

All authors conceived the original idea of the work. M.G.L. wrote the pipeline for training and evaluation of the models under the supervision of R.G and P.G. R.G performed the visual analysis of the embeddings. All authors discussed the results and contributed to the writing of the manuscript.

## Code and data availability

The full training and evaluation pipeline, including code and models, has been made publicly available on GitHub: https://github.com/edmanft/cat drug synergy under an Apache License 2.0. The original dataset can be accessed in the Supplementary Information section of Menden et al. [19].

## Declarations

### Ethics approval and consent to participate

Not applicable.

### Consent for publication

Not applicable.

