## Supplementary Information for "Optimizing drug synergy prediction through categorical embeddings in Deep Neural Networks"

### Supplementary information for
### Optimizing drug synergy prediction through categorical embeddings in Deep Neural Networks

Manuel González Lastre[1], Pablo González de Prado Salas[2] and Raúl Guantes[3,4,5,*]

[1]Departamento de Física Teórica de la Materia Condensada, Universidad Autónoma de Madrid, Spain
[2]Foqum Analytics, Spain
[3]Departamento de Física de la Materia Condensada, Universidad Autónoma de Madrid, Spain
[4]Materials Science Institute 'Nicolás Cabrera', Universidad Autónoma de Madrid, Spain
[5]Condensed Matter Physics Center (IFIMAC), Universidad Autónoma de Madrid, Spain

[*]

March 26, 2025

#### S1 Ablation analysis of random initialization

We performed an ablation analysis to evaluate the effect of random initialization on the performance of learned neural network embeddings across different machine learning models. In Table S1 we fixed the sklearn random seed and varied the embedding initialization, while in Table S2 we did the opposite, fixing the embedding initialization and varying the sklearn seed. We see that the results are consistent with the performance summary provided in the main text, where we vary both embedding and sklearn seeds simultaneously.

Aditionally, in Table S3 we report the improvement (or decline) in performance per model and embedding type against the best traditional encoding method.

Table S1: **Performance summary (leaderboard subset) of traditional encoding methods versus learned neural network embeddings across various models.** Results are reported as mean (standard deviation). Here we fix the sklearn seed and we change the embedding seed of the models.

| | Traditional encodings | | Neural Network encodings | | |
|---|---|---|---|---|---|
| **Model** | **LabelEncoder** | **OneHotEncoder** | **CategoryEmbedding** | **AutoInt** | **TabTransformer** |
| Linear Regression | 0.104 | **0.268** | 0.219 (0.021) | 0.249 (0.018) | 0.211 (0.021) |
| Ridge | 0.104 | 0.257 | 0.217 (0.025) | **0.260**(**0.018**) | 0.214 (0.016) |
| Lasso | 0.115 | 0.099 | **0.169**(**0.046**)* | 0.103 (0.017) | 0.099 |
| ElasticNet | 0.097 | 0.113 | **0.191**(**0.044**)* | 0.156 (0.017) | 0.105 |
| Bayesian Ridge | 0.090 | **0.268** | 0.216 (0.028) | 0.263 (0.015) | 0.223 (0.022) |
| SVR | 0.101 | 0.194 | 0.207 (0.041) | **0.256**(**0.009**)* | 0.222 (0.010) |
| GP | 0.112 | 0.118 | 0.016 (0.053) | **0.199**(**0.023**)* | 0.176 (0.011) |
| SGD | −0.008 | **0.273** | 0.221 (0.024) | 0.260 (0.017) | 0.223 (0.020) |
| Decision Tree | **0.152** | 0.098 | 0.119 (0.058) | 0.126 (0.036) | 0.101 (0.072) |
| Random Forest | **0.287** | 0.238 | **0.287**(**0.023**) | 0.280 (0.027) | 0.278 (0.037) |
| Extra Trees | 0.190 | 0.272 | **0.279**(**0.021**) | 0.267 (0.026) | 0.265 (0.035) |
| AdaBoost | 0.096 | 0.125 | 0.135 (0.034) | **0.193**(**0.034**)* | 0.164 (0.041) |
| Gradient Boosting | 0.205 | 0.258 | 0.224 (0.033) | **0.260**(**0.017**) | 0.250 (0.024) |
| XGB | 0.207 | 0.237 | **0.246**(**0.044**) | 0.233 (0.033) | 0.212 (0.042) |

An asterisk (*) indicates that the embedding method achieves at least 30% better performance compared to both Label Encoding (LE) and One-Hot Encoding (OHE).

Table S2: **Performance summary (leaderboard subset) of traditional encoding methods versus learned neural network embeddings across various models.** Results are reported as mean (standard deviation). Here we fix the embedding seed and we change the sklearn seed of the models.

| | Traditional encodings | | Neural Network encodings | | |
|---|---|---|---|---|---|
| **Model** | **LabelEncoder** | **OneHotEncoder** | **CategoryEmbedding** | **AutoInt** | **TabTransformer** |
| Linear | 0.104 | **0.268** | 0.179 | 0.239 | 0.190 |
| Ridge | 0.104 | 0.257 | 0.171 | **0.270** | 0.210 |
| Lasso | 0.115 | 0.099 | 0.134 | **0.144*** | 0.099 |
| ElasticNet | 0.097 | 0.113 | 0.122 | **0.171*** | 0.105 |
| Bayesian Ridge | 0.090 | **0.268** | 0.190 | 0.262 | 0.208 |
| SGD | −0.034 (0.091) | **0.268**(**0.004**) | 0.174 (0.015) | 0.257 (0.004) | 0.214 (0.004) |
| SVR | 0.101 | 0.194 | 0.135 | **0.262*** | 0.224 |
| GP | 0.112 | 0.118 | −0.049 | **0.220*** | 0.180 |
| Decision Tree | **0.146**(**0.018**) | 0.093 (0.029) | 0.039 (0.016) | 0.114 (0.018) | 0.123 (0.020) |
| Random Forest | 0.247 (0.017) | 0.256 (0.021) | 0.274 (0.014) | 0.269 (0.020) | **0.283**(**0.023**)* |
| Extra Trees | 0.205 (0.023) | 0.258 (0.011) | **0.259**(**0.016**) | 0.251 (0.017) | 0.261 (0.030) |
| AdaBoost | 0.089 (0.023) | 0.110 (0.018) | 0.138 (0.029) | **0.217**(**0.026**)* | 0.171 (0.028) |
| Gradient Boosting | 0.206 (0.005) | 0.257 (0.005) | 0.192 (0.009) | 0.245 (0.002) | **0.264**(**0.004**) |
| XGB | 0.207 | **0.237** | 0.219 | 0.213 | 0.159 |

An asterisk (*) indicates that the embedding method achieves at least 30% better performance compared to both Label Encoding and OneHot Encoding.

Table S3: **Percentage of improvement of learned neural network embeddings over traditional encoding methods across various models.** Values represent the percentage of improvement (or decline) compared to the best of LabelEncoder and OneHotEncoder. Net improvement shows the average percentage improvement across all models.

| Model | AutoInt | CategoryEmbedding | TabTransformer |
|---|---|---|---|
| Linear | −7.00 | −18.24 | −21.23 |
| Ridge | 1.17 | −15.59 | −16.77 |
| Lasso | −10.21 | **46.88** | −14.14 |
| ElasticNet | 38.55 | **69.59** | −6.52 |
| Bayesian Ridge | −2.10 | −19.38 | −16.83 |
| SGD | −3.29 | −20.13 | −15.75 |
| SVR | **31.86** | 6.78 | 14.56 |
| GP | **68.39** | −86.61 | 48.84 |
| Decision Tree | −22.36 | −17.57 | −35.76 |
| Random Forest | 10.84 | 7.25 | **13.99** |
| Extra Trees | 4.41 | **8.61** | 7.81 |
| AdaBoost | **87.80** | 16.01 | 31.87 |
| Gradient Boosting | 1.09 | −14.97 | −2.74 |
| XGB | −1.43 | 3.99 | −10.55 |
| **Net Improvement** | **17.85** | −24.95 | −1.47 |

#### S2 Embedding analysis

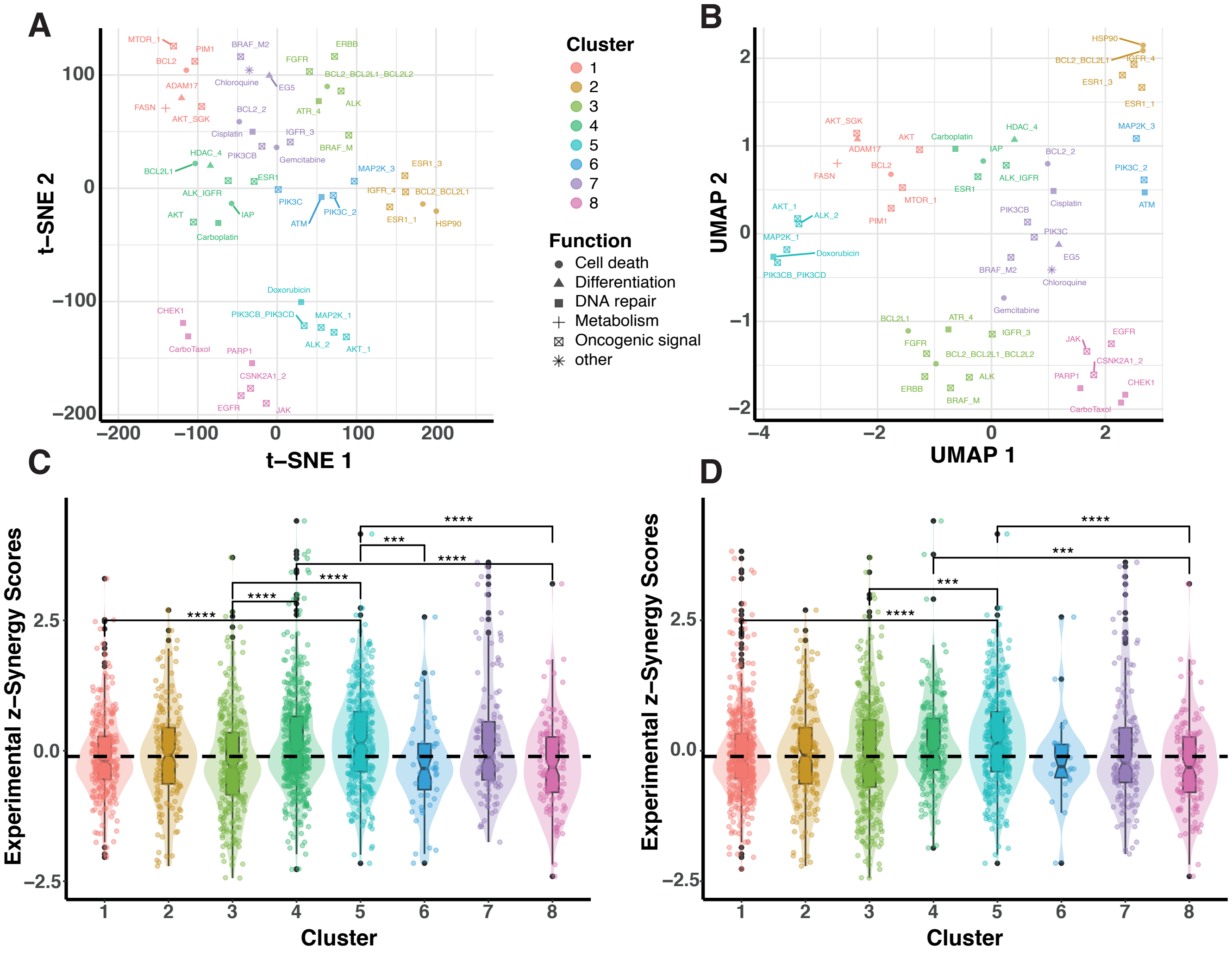


Figure S1: **Relationship between embeddings revealed by dimensionality reduction are robust.** Analysis of embedding vectors of the categorical variable `Compound B`, for the best performing `CategoryEmbedding` training run. **A.** Embedding projections on the first two principal dimensions of t-distributed stochastic network embedding(t-SNE), see Section 2.6 in main text. **B.** Embedding projections on the first two principal dimensions of Uniform Manifold Approximation and Projection (UMAP). The adjusted rand index between clusters in panels **B, C** is 0.77. Overlap between both clusterings is complete except for Clusters 6 and 7 that differ in one element. **C.** Boxplots with overlayed distributions (violin plots) of the z-normalized experimental synergy scores corresponding to the drug combinations containing `Compound B`, grouped by the clusters shown in panel **A**. Shown with bars are the most significant differences between distribution means. Stars represent adjusted (Bonferroni) p values calculated with two-sided Wilcoxon rank-sum tests (ns: $p > 0.05$; *: $p \leq 0.05$, **: $p \leq 0.01$; ***: $p \leq 0.001$; ****: $p \leq 0.0001$). The dashed horizontal line represents the median value of the whole distribution of synergy scores. **D.** Boxplots of experimental synergy scores corresponding to the clusters in UMAP two-dimensional projection shown in panel **B**

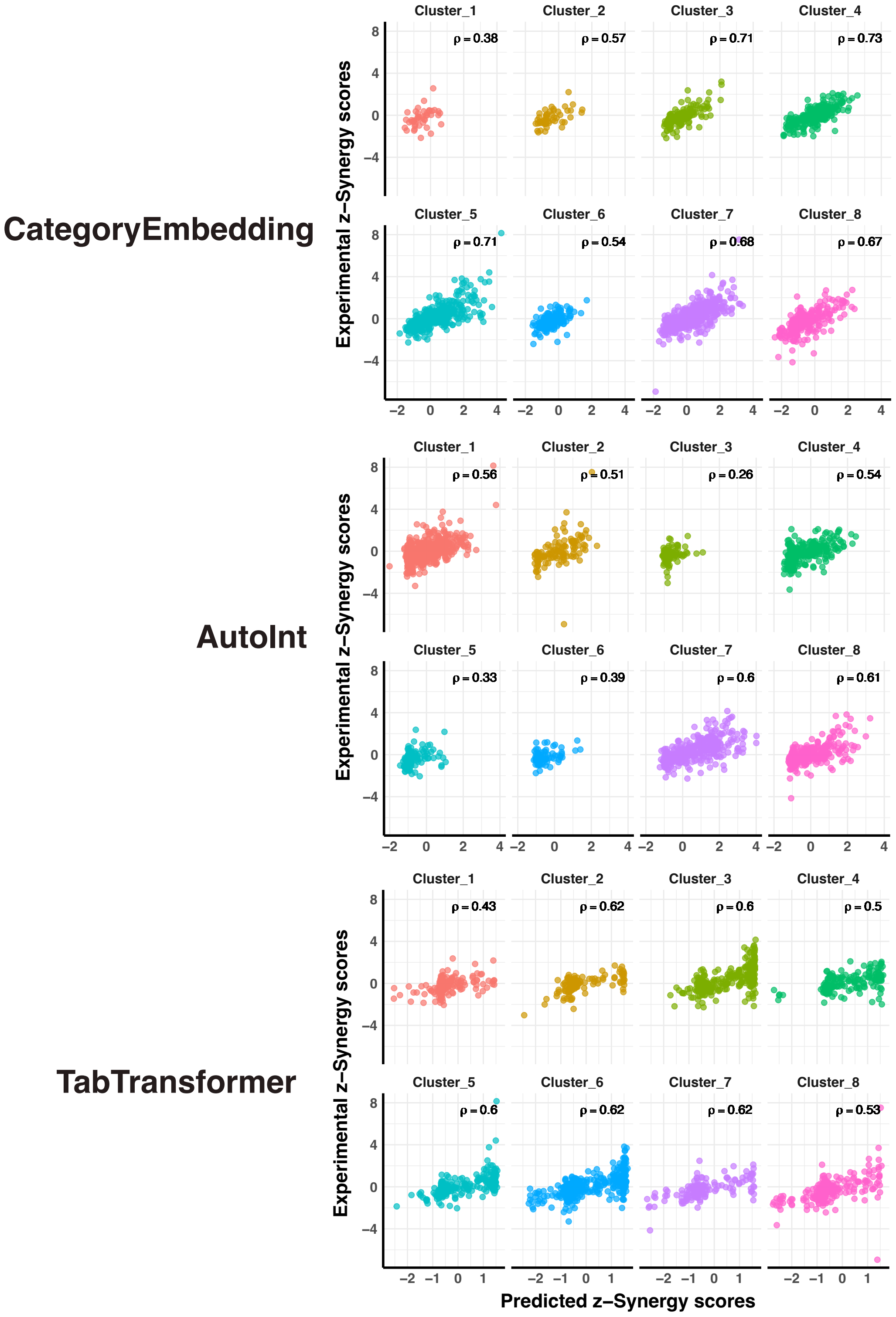


Figure S2: **Correlation between experimental and predicted synergy scores for different embedding methods.** Scatter plots and corresponding Pearson correlation coefficients ($\rho$) for the clusters shown in Figure 3 in main text. The overall Pearson correlation coefficients are $\rho = 0.7$ (CategoryEmbedding), $\rho = 0.59$ (AutoInt) and $\rho = 0.61$ (TabTransformer).

#### S3 Ensembling for competitive performance

We explore two simple ensembling strategies for models trained on the embeddings of a `CategoryEmbedding` model with a respectable performance WPC = 0.282 (seed 42):

- **Greedy forward selection with uniform averaging**, where models are added sequentially based on the improvement in weighted Pearson correlation (WPC) when their predictions are uniformly averaged with the current ensemble.

- **Greedy forward selection with weighted averaging**, where at each step, we optimize a set of weights over the selected models to maximize WPC.

All ensemble selection procedures are run until the performance saturates ($\Delta$WPC = 0). For the uniform averaging ensembling (Table S4), we achieve a final WPC = 0.319 by averaging the predictions of 7 different models, including tree-based methods (Random Forest, Extra Trees, Gradient Boosting), linear models (Lasso, Bayesian Ridge), and kernel-based models (SVR). This diversity reflects the benefit of combining models with different inductive biases, each capturing distinct aspects of the feature space encoded by the learned embeddings.

The weighted ensemble (Table S5), while only marginally better in terms of WPC (0.327 vs. 0.319), is more compact, selecting only four models. The optimization of weights seems to effectively prune redundant or low-value models, resulting in a more efficient ensemble with fewer components but improved synergy.

Notably, these results approach the average experimental performance defined by biological replicates (WPC $\sim$ 0.4) [1], despite being trained on a minimal set of features—only categorical descriptors and monotherapy response curves. This highlights the strength of the learned embeddings in capturing relevant pharmacological and cellular information. In contrast to many approaches in the original DREAM Challenge, which relied heavily on molecular and genomic profiling, our ensemble reaches competitive performance using low-dimensional and easily accessible data.

Table S4: **Greedy forward selection with uniform averaging.** At each step, models are added based on improvement in ensemble WPC.

| **Step** | **Model Added** | **WPC** | **ΔWPC** |
|---|---|---|---|
| 1 | Random Forest | 0.290 | — |
| 2 | Stochastic Gradient Descent | 0.302 | 0.012 |
| 3 | Extra Trees | 0.307 | 0.006 |
| 4 | Support Vector Regression | 0.310 | 0.003 |
| 5 | Bayesian Ridge Regression | 0.313 | 0.002 |
| 6 | Gradient Boosting | 0.314 | 0.001 |
| 7 | Lasso Regression | 0.319 | 0.006 |
| 8 | Gaussian Process | 0.319 | 0.000 |

Table S5: **Greedy forward selection with weighted averaging.** At each step, models are added based on improvement in ensemble WPC.

| Step | Model Added | WPC | ΔWPC |
|---|---|---|---|
| 1 | Random Forest | 0.290 | — |
| 2 | ElasticNet Regression | 0.309 | 0.018 |
| 3 | Stochastic Gradient Descent | 0.320 | 0.011 |
| 4 | Gradient Boosting | 0.327 | 0.007 |
| 5 | XGBRegressor | 0.327 | 0.000 |
